# Analysis Of Molecular Networks In The Cerebellum In Chronic Schizophrenia: Modulation By Early Postnatal Life Stressors In Murine Models

**DOI:** 10.1101/2021.02.21.432145

**Authors:** América Vera-Montecinos, Ricard Rodríguez-Mias, Karina S. MacDowell, Borja García-Bueno, Juan C. Leza, Judit Villén, Belén Ramos

## Abstract

Despite the growing importance of the cortico-cerebellar-thalamo-cortical circuit in schizophrenia, limited information is available regarding altered molecular networks in cerebellum. To identify altered protein networks, we conducted proteomic analysis of grey matter of postmortem cerebellar cortex in chronic schizophrenia subjects (n=12) and healthy individuals (n=14) followed by an extensive bioinformatic analysis. Two double-hit postnatal stress murine models for SZ were used to validate the most robust candidates. The models were maternal deprivation combined with an additional stressor: social isolation or chronic restraint stress. We found that the individual proteomic profile allowed the segregation of most schizophrenia cases from healthy individuals. We found 250 proteins with altered levels. This group was enriched in proteins related to mental disorders, mitochondrial disease, stress, and a number of biological functions including energy, immune response, axonal cytoskeletal organization and vesicle-mediated transport. Network analysis identified three modules: energy metabolism, neutrophil degranulation and a mixed module of mainly axonal-related functions. We analysed the most robust candidates in the networks in two double-hit stress murine models. METTL7A from the degranulation pathway was reduced in both models, while NDUFB9 from the energy network and CLASP1 from the axonal module decreased in only one model. This work provides evidence for altered energy, immune and axonal-related networks in the cerebellum in schizophrenia, suggesting that the accumulation of molecular errors, some by an early postnatal stress exposure, could lead to a failure in the normal cerebellar functions, impairing synaptic response and the defence mechanisms of this region against external harmful injuries in schizophrenia.

## INTRODUCTION

Schizophrenia (SZ) is a complex polygenic psychiatric disorder involving dysregulation of multiples pathways [1] with an estimated prevalence up to 1% in the general population [2] and a high heritability up to 79% [3]. Although the aetiology of SZ is not fully understood, several hypotheses have been postulated. The neurodevelopmental hypothesis proposes two critical periods of neurodevelopmental vulnerability, early life and adolescence. An environmental double-hit in these phases in a genetically predisposed individual is required for the emergence of the disease [4,5]. Based on this hypothesis, cumulative damage in different molecular pathways required for the early development of the central nervous system (CNS) could contribute to the failure of axonal assembly connections and normal synaptic transmission, which could remain latent until adolescence [6]. In this phase of life, an additional stressor such as psychosocial stress could impact on these vulnerable pathologic neural circuits leading to altered functioning of synaptic responses and the emergence of symptoms [7–9]. To study the role that psychosocial stress may play in SZ, different animal models have been developed that include prenatal or perinatal stress. Prenatal models may include different types of stressors such as restraint of movement together with water and/or food deprivation [9] and foot-shocks [10] to the mother. Supporting these animal models are other findings from animal experiments that showed that prenatal exposure to stressors leads to learning deficits [11,12]. Furthermore, a study found that post-weaning social isolation induces altered adult behaviour as a result of hyperactivity of the hypothalamic-pituitary-adrenal axis [13]. Another animal model used to understand the origin of SZ is the double-hit model in which two hits are required for the emergence of this disorder, the first hit occurring in the prenatal or perinatal phase and a second during adolescence [14,15]. SZ-like symptoms are reported in these models. Adolescence is an active phase of remodelling of neuronal networks and synaptic activity in which significant energy demand is required for the correct building of functional circuits [16]. Thus, injuries in this period could compromise this active phase of remodelling of circuits, altering their functioning and promoting disease emergence.

In the last decades, the cortico-cerebellar-thalamo-cortical circuit (CCTC) has been proposed to play a key role in cognitive impairments and symptoms in individuals at ultra-high risk for SZ and in individuals that have transited to SZ [17–19]. The cerebellum is a brain area that forms part of this circuit that modulates synaptic responses of cortical regions. This area-has been proposed to play an important role in SZ pathophysiology [17,20,21]. Nowadays, clinical and neuroimaging studies suggest that the cerebellum supports cognitive and executive functions in SZ [22–24]. The cerebellum integrates input signals from the cortical areas to generate output signals back to the same area in the cortex. This is important to detect and correct errors during the execution of motor movements that it could be also implicated in the regulation of cognitive functions [25]. The cerebellum is the last region to complete neuronal progenitor division, neuronal migration, and pruning of dendritic arborization [26]. This lengthy maturation phase of the cerebellum makes this area highly vulnerable to error accumulation during development, which may have a significant impact in SZ. Thus, molecular alterations in the cerebellum in SZ constitute an attractive biological substrate as a possible reservoir of failures in multiple pathways through development and during the disease.

Despite the important role of the cerebellum in SZ, only a few recent studies have investigated global mRNA [27] or proteomic alterations in this region [28,29]. The main aim of our study was to identify altered molecular networks in the cerebellum of SZ patients. To achieve this, we compared the proteomic profile in *postmortem* lateral cerebellar cortex of individuals with chronic schizophrenia (n=12) and control healthy subjects (n=14), that we obtained using single shot liquid chromatography-tandem mass spectrometry analysis (Figure 1A).

**Figure 1.**
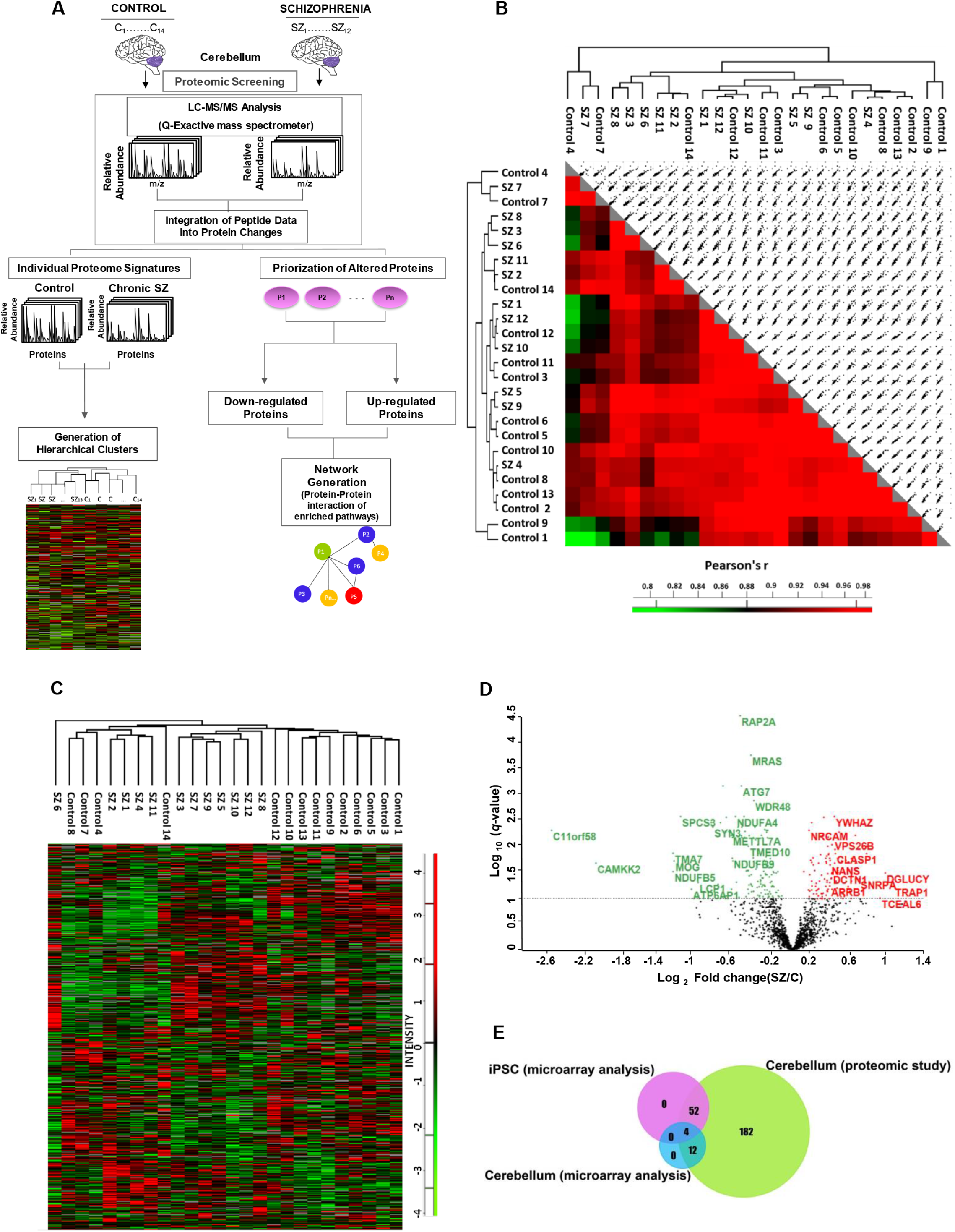
Quantitative proteomic analysis in the cerebellum in chronic schizophrenia. **A.** Experimental design for proteomic analysis to identify altered pathways in schizophrenia. Protein lysates from the *postmortem* cerebellum of control (C) (n=14) and chronic schizophrenia (SZ) patients (n=12) were processed as described. The peptides were separated and analysed by liquid chromatography (LC) coupled to tandem mass spectrometry. The relative fold change of peptides was integrated into protein changes. The individual protein signatures for each case and control were used to generate hierarchical clusters. The prioritization of altered proteins (P1-Pn represents generic proteins) in SZ was obtained by comparing protein fold changes between control and SZ groups (significant proteins adjusted using an FDR of 0.1). We performed two analyses for these altered proteins in SZ: **i**) Unsupervised hierarchical clustering analysis generated from quantified proteins in 12 SZ and 14 healthy control samples of *postmortem* cerebellum; **ii**) Generation of networks from significantly enriched pathways by protein-protein interaction. These analyses were performed using Perseus and String, respectively. **B.** A correlation matrix for 1474 quantified proteins across sample pairs. **C.** Unsupervised hierarchical clustering analysis was obtained with matrix processing according to the Euclidean distance and z-score aggregation method. Protein profiles were generated from 1474 quantified proteins in 12 SZ and 14 healthy control samples of *postmortem* cerebellum and were clustered according to the z-score and displayed as a heat map. Green colour clusters represent underexpressed proteins. Red colour clusters represent overexpressed proteins. **D.** Volcano plot of the −log_10_ *q*-value (adjusted *p*-value; FDR (≤ 0.1)) *versus* the log_2_ fold change in the cerebellum in SZ relative to healthy control. SZ, schizophrenia; C, control. Up-regulated and down-regulated significant proteins are represented in red and green, respectively. The grey line shows the FDR threshold. **E.** Venn diagram showing overlap between proteins previously reported in SZ through gene expression analysis obtained from SZDB (human cerebellum and iPSC) and our proteomic study in cerebellum.

Furthermore, we validated robustly altered proteins on two separate double-hit murine models of SZ induced by maternal deprivation combined with an additional stressor (social isolation or chronic restraint stress).

## MATERIALS AND METHODS

### *Postmortem* human brain tissue

Samples from the cerebella of subjects with chronic schizophrenia (n=12) and healthy controls (n=14) were obtained from the neurologic tissue collection of the *Parc Sanitari Sant Joan de Déu* Brain Bank[30] and the Institute of Neuropathology of the *Universitari de Bellvitge* Hospital respectively. All SZ subjects were institutionalized donors with long-term illness who had no history of neurological episodes. We matched SZ and control groups by gender (only male patients were included), age, *postmortem* delay (PMD) and pH (Table S1). Experienced clinical examiners interviewed each donor *antemortem* to confirm SZ diagnosis according to the Diagnostic and Statistical Manual of Mental Disorders (DSM-IV) and International Classification of Diseases 10. All deaths were due to natural causes. The study was approved by the Institutional Ethics Committee of Parc Sanitari Sant Joan de Déu. A written informed consent was obtained from each subject. The last daily chlorpromazine equivalent dose for the antipsychotic treatment of patients was calculated based on the electronic records of the last drug prescriptions administered up to death as described previously [31]. Human cerebellar lateral cortex was dissected from coronal slabs stored at −80°C, extending from the pial surface to white matter only including grey matter.

### Label free quantification proteomic analysis mass spectrometry

Protein extracts were prepared as described previously [32]. Protein concentration was determined by Bradford assay (Biorad, Hercules, CA, USA). Proteomic samples were prepared as described in the supplementary methods, using a starting protein amount of 200 μg Peptide mixtures were analyzed by single-shot liquid chromatography-tandem mass spectrometry (LC-MS/MS) in a Q-Exactive mass spectrometer (Thermofisher Scientific, CA, USA) using a top 20 data-dependent acquisition method. Mass spectrometry data was analyzed using MaxQuant. The target-decoy database search strategy was used to guide filtering and estimate false discovery rates (FDR). Peptides matches were filtered to an FDR of ≤0.01. The minimum required peptide length was seven residues. Label-free quantification (LFQ) was selected for individual protein comparisons between control and SZ groups. Proteins quantified in fewer than 7 samples per group were excluded from the analysis. The normalized LFQ intensity was referred to the mean intensity of the controls. A significance value for each quantified protein was calculated using the Student’s t-test and adjusted for multiple-hypothesis testing using the Benjamini-Hochberg method [33]. FDRs were computed for all significant values and the FDR threshold was set to 0.1. The quantified proteins were imported into the Perseus software platform (version 1.6.1.3) for quality control and further analysis [34]. See supplementary methods for more detailed information.

### Bioinformatic analysis

To visually identify significantly altered proteins, we plotted the log_2_ of the fold change of normalized LFQ intensities between schizophrenia and control samples along with the FDR adjusted −log_10_ (*q*-value).

We used the Schizophrenia Database [35] and FunRich Tool v.3.1.3 [36] to compare the identified altered proteins to those previously reported in gene expression studies. To perform the disease, gene ontology (GO), and pathway analyses we used the Webgestalt-mediated ORA method. Disease terms were obtained from the PharmacoGenetics Knowledge Base (PharmGKB) and Gene List Automatically Derived For You (GLAD4U) databases [37,38]. For the GO analysis, we performed a non-redundant enriched categories analysis. For the pathway analysis, we used the Reactome database, and for the network generation, we used String version 11.0 [39]. To perform the screening on protein localization analysis in different tissue layers in the cerebellum we used the Human Protein Atlas database [40]. The enrichment analyses were set to an FDR of 0.1.

### Murine models

Three pregnant Wistar rats (Harlan Ibérica, Spain) at gestation day 15 were individually housed in a controlled environment in a 12h light/dark cycle with free access to food and water. One of the litters was used as a control group and the other as a double-hit model randomly. After birth, at postnatal day (PD) 9, both litters were exposed to maternal deprivation for 24hrs as a first hit. On PD21, the pups were weaned and one of the litters was isolated for 5 weeks as a second hit (PD21-56); the pups were housed individually and denied physical contact with their siblings, although they could smell, hear and see them. After isolation, the pups were regrouped (MD/Iso). Meanwhile, the other litter was exposed to restraint stress between PD 72 and 78 for 6hr every day (MD/RS). These conditions represent well suited murine models for the study of neuropsychiatric dysfunctions [41,42]. Group sample sizes were CT, n=11; MD/Iso, n=9; MD/RS, n=7. All experimental protocols adhered to the guidelines of the Animal Welfare Committee of the Complutense University in accordance with European legislation (D2010/63/UE). The animals were subjected to cervical dislocation. The brain was removed from the skull and the cerebellum was excised and frozen at −80°C until assayed. Samples were homogenized by sonication in PBS (pH=7) mixed with a protease inhibitor cocktail (Complete^®^, Roche, Spain). After adjusting protein levels, homogenates of cerebellar tissue were mixed with loading buffer and 15μg were loaded into an electrophoresis gel, then blotted onto a nitrocellulose membrane with a semi-dry transfer system (Bio-Rad) and incubated with specific antibodies against: (1) Methyltransferase-like 7A (METTL7A); (2) NADH dehydrogenase [ubiquinone] 1 beta subcomplex subunit 9 (NDUFB9); (3) CLIP-associating protein 1 (CLASP1); (4) 14-3-3 protein zeta/delta (YWHAZ); (5) Beta-actin (β-actin). See supplementary methods for more detailed information.

### Statistical analysis

Normal distribution of variables was determined using the D’Agostino-Pearson test. Demographics and tissue features of the samples were compared between cases and controls using the Student’s *t*-test for parametric quantitative variables and the Mann-Whitney U test for non-parametric variables. Spearman or Pearson correlation analysis was carried out to detect association of our proteomic data with other clinical-, demographic- and tissue-related variables (age, *postmortem* delay, pH, daily chlorpromazine equivalent dose, and duration of illness). Statistical analysis was performed with Graph Prism version 7.00. The significance level was set to 0.05.

## RESULTS

### Quantitative proteomic analyses in cerebellum in chronic schizophrenia

To identify altered proteins related to SZ, we performed a proteomic analysis of human cerebellar lateral cortex protein extracts from 12 male SZ patients and 14 control individuals matched for gender, age and *postmortem* delay. No differences were observed between the SZ and control groups for any demographic or tissue-related variables (Table S1). Using mass spectrometry, a total of 2578 proteins were quantified. 1474 proteins (57%) were quantified in at least 7 individuals per group and used for subsequent bioinformatics analyses (Supplementary data 1). 97.8% of the proteins were identified with 2 or more peptides and 46.5% with five or more peptides. For the individual proteome signature analyses, we examined the similarity of the individual proteome through a correlation matrix. To assess the similarity between the proteomes of the different individuals, we calculated Pearson correlation coefficients and visualized the results in a correlation matrix (Figure 1B). All correlations were above 0.7 (Figure 1B). To assess the similarity between the SZ and control groups we calculated the Pearson correlation coefficient of the protein intensity means calculated for each group. This correlation was 0.989, indicating that globally the cerebellar proteomes of SZ and control individuals were highly similar (Data not shown). Unsupervised hierarchical clustering analysis of the proteomic profiles was able to segregate controls and SZ samples, with the exception of five controls (#8, #7, #4, #12 and #14) (Figure 1C). We further investigated whether the demographic- and tissue-related features could explain the differential clustering of these controls and observed that although four of these controls segregated together, none of the variables influence this segregation (Figure S1). We identified 250 proteins significantly regulated (16.9%) in SZ with an FDR of <0.1 (Supplementary dataset 2) (142 proteins down-regulated and 108 up-regulated). None of the regulated proteins showed significant correlation with PMD or pH (FDR<0.1) (Supplementary datasets 3 and 4) was found for any altered protein. A volcano plot was used to categorize proteins as up or down-regulated based on the fold change (log_2_ FC) between SZ and control cases and the corrected *p*-value (−log_10_ (*q*-value) adjusted to FDR<0.1). This analysis revealed a range of changes, where most of the significant changes displayed between 0.2 and 0.6 Log_2_ FC (Figure 1D). We found that only 56 of the 250 altered proteins had been previously reported in a gene expression study in an iPSC model of SZ, and that only 16 altered proteins had been reported in a microarray assay in human cerebellum (Figure 1E, Supplementary dataset 2). Additionally, 86 altered proteins in the cerebellum had been previously reported as altered in other brain regions in SZ and 20 proteins have been reported in two recent cerebellar proteomic studies (Table S1 and Supplementary dataset 2).

### Gene ontology enrichment analysis

The gene ontology (GO) enrichment analysis showed enrichment of disease categories related to mitochondrial diseases and mental disorders for down-regulated proteins, but in stress, neurodegenerative diseases, and drug interaction with drug for up-regulated proteins (Figure 2A and Table 1). The analysis of biological processes of the 142 down-regulated proteins revealed significant enrichment of terms related to energy metabolism (Figure 2B and Table 1). For the 108 up-regulated proteins, the enriched categories were related to structural and signalling functions (Figure 2B and Table 1). For the down-regulated proteins, two predominant pathways were enriched: the citric acid (TCA) cycle/respiratory electron transport and neutrophil degranulation (Figure 2C). For the up-regulated proteins, six pathways were enriched. From the highest to lowest score of significance, these were: vesicle-mediated transport, apoptosis, Rho GTPase effectors, signalling by Rho GTPases, axon guidance, and cell cycle (Figure 2C).

**Figure 2.**
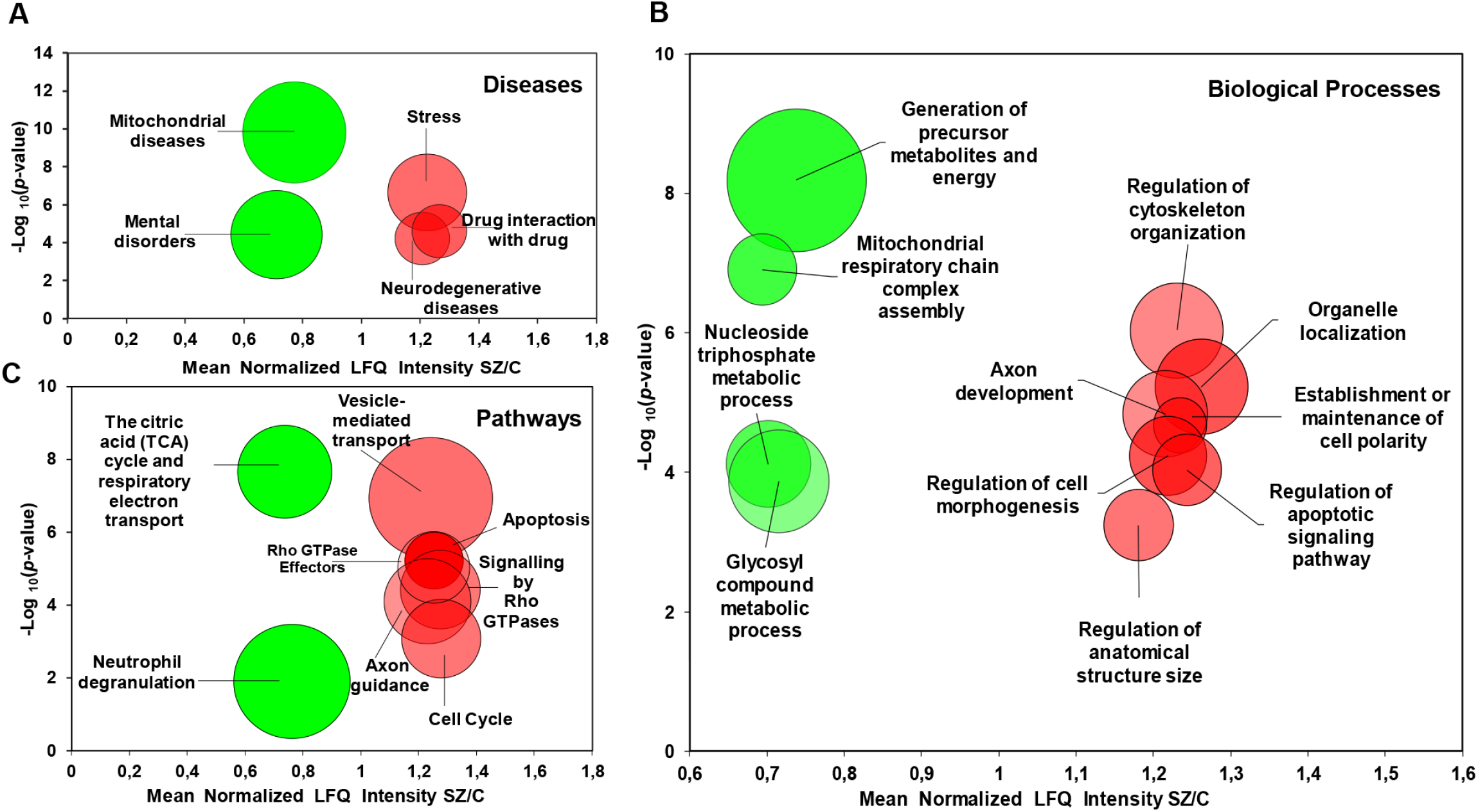
Enrichment analyses from proteome cerebellum in chronic schizophrenia. **A.** The bubble chart showing enriched disease categories (A), biological processes (B) and pathways (C) for 142 down-regulated proteins and 108 up-regulated proteins in SZ. **A**. The enriched categories for the down-regulated proteins were: mitochondrial diseases (PA447172), mental disorders (PA447208), and for the up-regulated proteins: stress (PA445752), drug interaction with drug (PA165108622), and neurodegenerative diseases (PA446858). **B.** Non-redundant enriched biological processes categories for down-regulated proteins in SZ were: generation of precursor metabolites and energy (GO: 0006091), mitochondrial respiratory chain complex assembly (GO: 0033108), nucleoside triphosphate metabolic process (GO: 0009141), glycosyl compound metabolic process (GO: 1901657). For up-regulated proteins, the enriched functions were: regulation of cytoskeleton organization (GO: 0051493), organelle localization (GO: 0051640), axon development (GO: 0061564), establishment or maintenance of cell polarity (GO:0007163), regulation of cell morphogenesis (GO:0022604), regulation of apoptotic signalling pathway (GO: 2001233), regulation of anatomical structure size (GO: 0090066), and microtubule-based movement (GO: 0007018). **C.** The enriched pathway categories in SZ for the down-regulated proteins were: citric acid (TCA) cycle/respiratory electron transport (R-HSA-1428517) and neutrophil degranulation (R-HSA-6798695). The enriched pathways for the up-regulated proteins were: vesicle-mediated transport (R-HSA-5653656), apoptosis (R-HSA-109581), signalling mediated by Rho GTPase effectors (R-HSA-195258), signalling by Rho GTPases (R-HSA-194315), axon guidance (R-HSA-422475), and cell cycle (R-HSA-1640170). The X-axes show the mean of normalized LFQ intensity in SZ relative to control group for all the proteins that belonged to each category. The Y-axes show the −log_10_ enrichment *p*-value. The bubble size is directly proportional to the number of proteins represented in each enriched category of diseases, biological processes or pathways. Red colour represents up-regulated proteins. Green colour represents down-regulated proteins. SZ, schizophrenia; C, control.

**Table 1:**
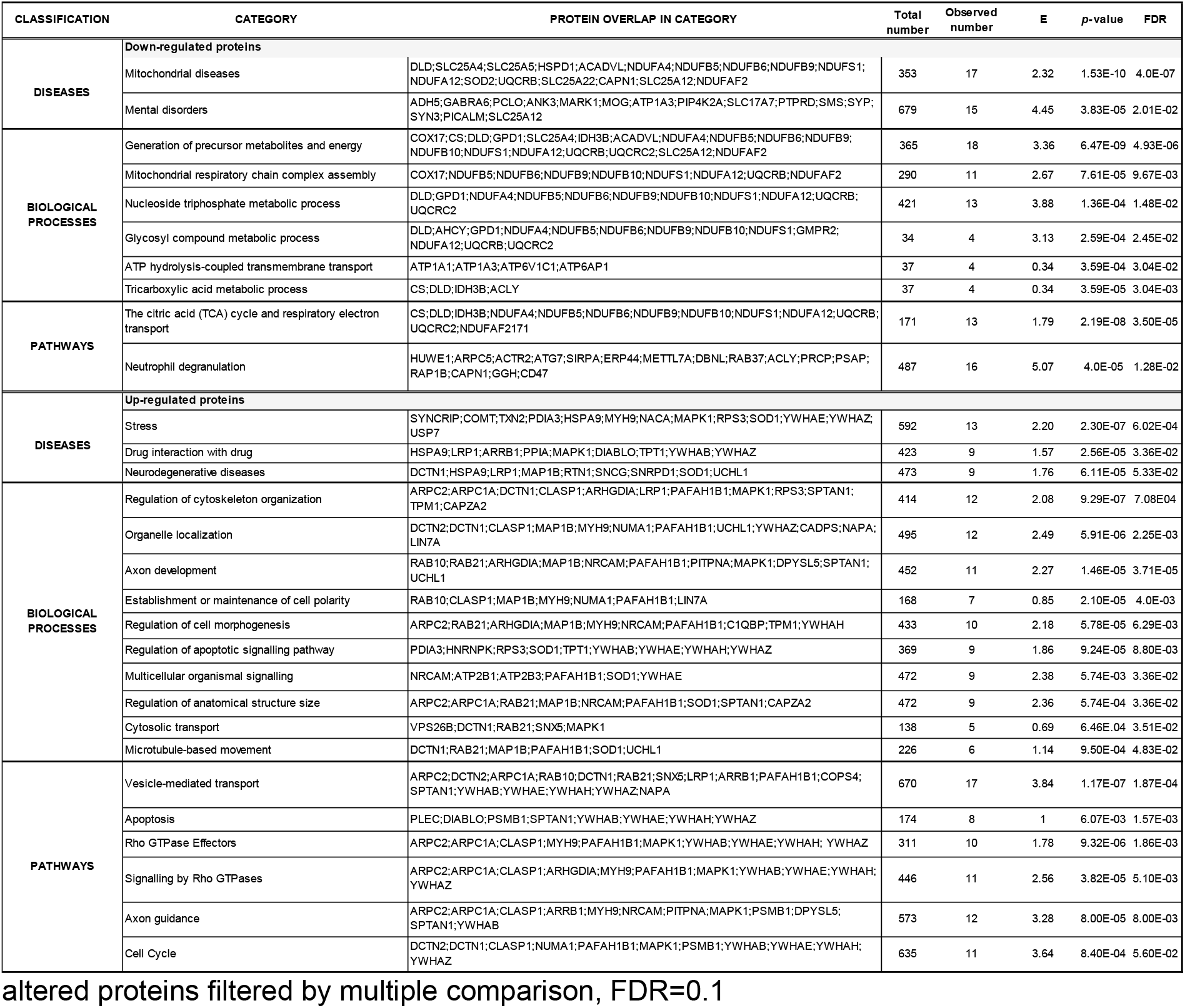
Non-redundant categories of disease, gene ontology and pathways among 250

### Network generation from enriched pathways in the cerebellum

Network analysis of enriched pathways for the down-regulated proteins revealed two distinct modules, one related to energy metabolism with TCA cycle/respiratory electron transport proteins, and another consisting of neutrophil degranulation pathways (Figure 3A). The neutrophil degranulation proteins overlapped with secretory granular and secretory lumen pathways. For the enriched up-regulated pathways, the network analysis showed overlapping pathways that were mainly related to axonal development and functioning (Figure 3B). The vesicle-mediated transport pathway showed the highest overlap with other pathways, with 52% overlapping with signalling by Rho GTPases proteins, 35% with the cell cycle, 29% with apoptosis, and 23% with axon guidance proteins (Figure 3B). Next, for the proteins in the aforementioned modules, we used publicly available immunocytochemistry data to assess their expression in the different layers of the cerebellum and found that all the proteins were widely distributed throughout the cerebellar layers (Figure 3) with the exception of proteins: ARRB1, ATG7 and SNX5 which were only expressed in Purkinje cells, and CD47 which was only expressed in the granular layer. For each network module, we identified one or two robust protein candidates for further characterization, based on LFQ-intensity fold change and coefficient of variation (<0.35) (Figure 3; Supplementary data 5). The selected proteins were NDUFB9 from the energy module, METTL7A from the neutrophil degranulation module, and YWHAZ (vesicle-mediated transport) and CLASP1 (axonal guidance) for the mixed module (Figure 3; Supplementary data 5).

**Figure 3.**
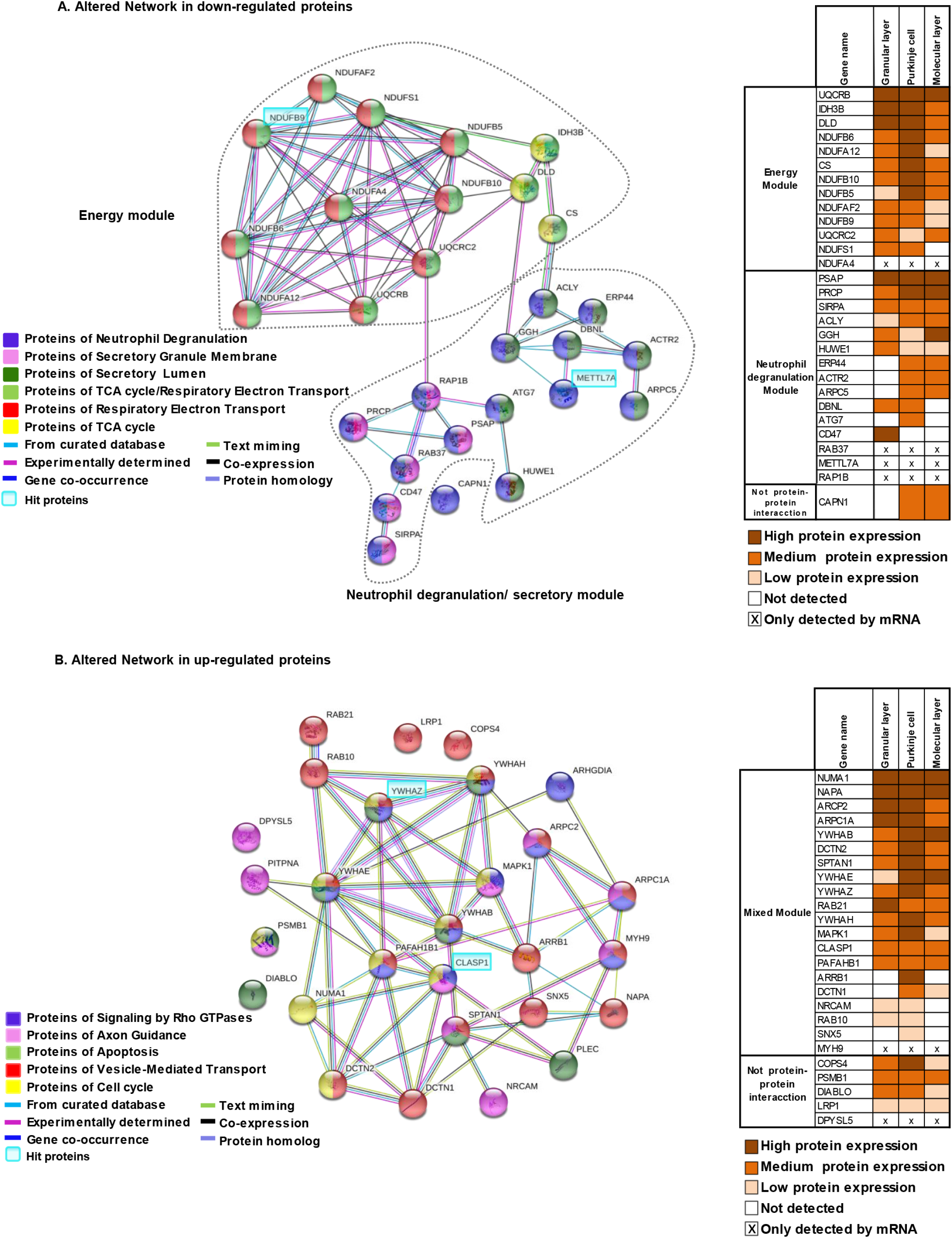
Network generation formed by altered pathways in cerebellum. **A.** A protein-protein interaction network illustrates the interaction and overlap between proteins of the down-regulated pathways. Two modules were obtained: an energy metabolism module and a neutrophil degranulation/secretory module. **B.** A protein-protein interaction network was generated using the up-regulated pathways. The interaction overview shows how proteins overlap in the different pathways. The right-hand panels show the level of protein expression determined by immunohistochemistry for each protein in the down- and up-regulated modules and their localization in the different layers in the cerebellum. These pathways were significantly enriched in the panel of 250 altered proteins in chronic SZ cerebellum. Each node represents a protein. Colour denotes membership to the module. The coloured edge (connections between nodes) represents the type of interaction between nodes. Highlighted gene symbols represent the most robust hit protein for each module based on the most prominent fold change with a coefficient of variation lower than 0.35 in both control and SZ groups. In the mixed module, the candidate selected belonged to at least three different pathways. Both networks were generated using the String V.11. The Human Protein Atlas database was used for the levels of protein expression determined by immunohistochemistry. Protein expression localization in the right panels was obtained from the Human Protein Atlas database.

### Validation of altered pathways in double-hit SZ murine models

We further investigated significantly regulated proteins NDUFB9, METTL7A, CLASP1 and YWHAZ in the cerebellum of two independent double-hit murine models for SZ using Western blot. The models were maternal deprivation combined with an additional stressor: social isolation (MD/Iso) or chronic restraint stress (MD/RS) (Figure 4). We observed that NDUFB9 levels were reduced in the human SZ cohort and in the MD/Iso model but not in the MD/RS model (Figure 4A and Figure S2A). METTL7A showed a decrease in protein expression levels in the chronic SZ cohort and in both murine models (Figure 4B and Figure S2B). CLASP1 was down-regulated in the MD/Iso model but up-regulated in the SZ cohort (Figure 4C and Figure S2C). No significant changes were found in the DM/RS model for CLASP1. YWHAZ, which was up-regulated in the SZ cohort, did not show any change in either of the two SZ murine models (Figure 4D and Figure S2D). Thus, METTL7A was the only one that was consistently deregulated in the human SZ cohort and the two murine models.

**Figure 4.**
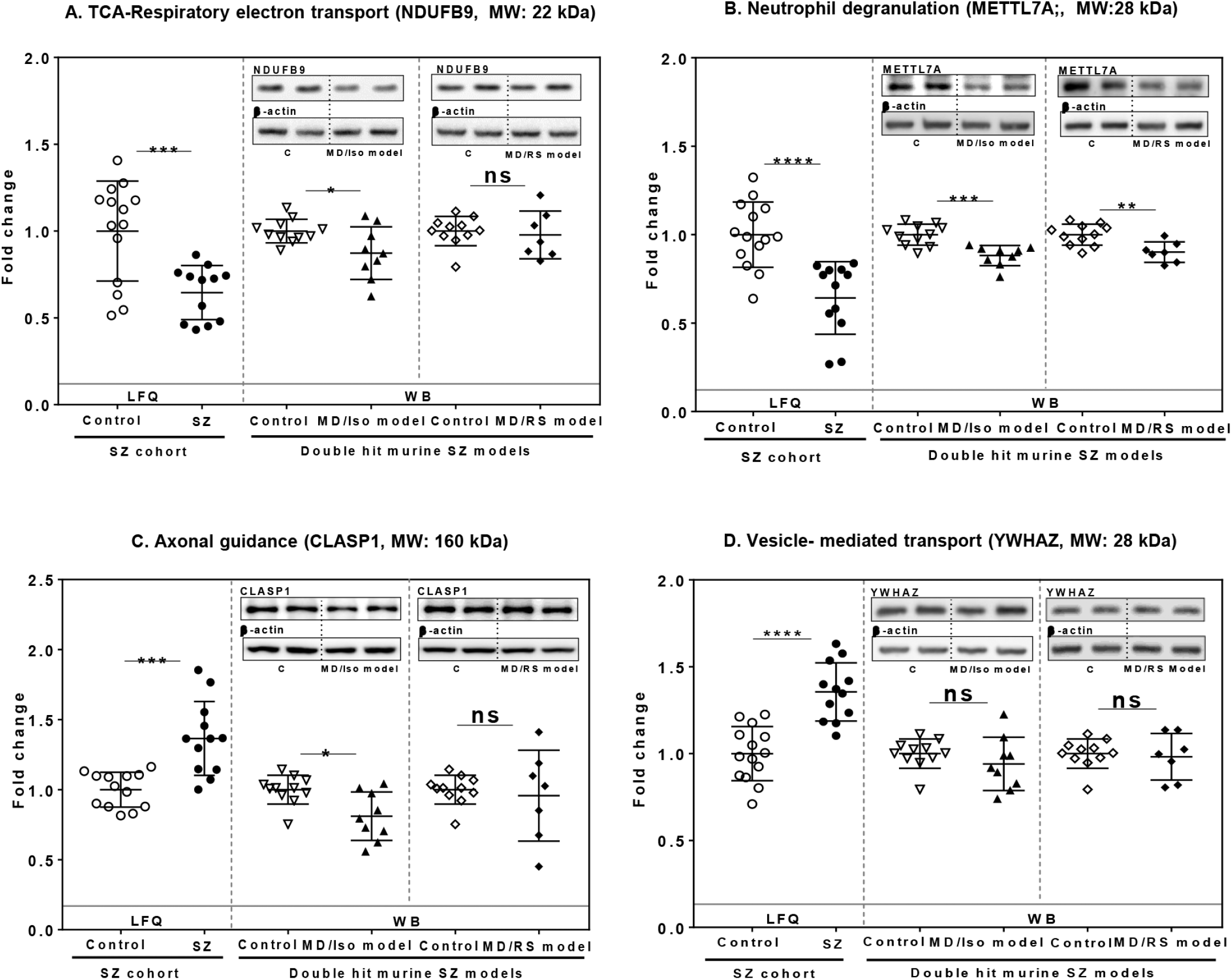
Analysis of hit proteins from altered pathways in a human SZ cohort and two double-hit SZ murine models. Protein levels of candidate hit proteins METTL7A (**A**), NDUFB9 (**B**), CLASP1 (**C**) and YWHAZ (**D**) from the indicated enriched pathway in SZ were analysed in the cerebellum of the human SZ cohort compared to the two double-hit SZ murine models, maternal deprivation combined with social isolation (MD/Iso) or chronic restraint stress (MD/RS). The protein fold change in the human SZ cohort (control: n=14; SZ=12) was determined by proteomic analysis as explained in the methods and the relative fold change in the murine models was determined by immunoblot analysis (control: n=11; MD/Iso model: n=9; MD/RS model: n=8). TCA-RET: citric acid cycle-respiratory electron transport. VTM: vesicle-mediated transport. Statistical analysis was performed using the Student’s t-test for samples with normal distribution and the Mann-Whitney U test was carried out for non-parametric distribution in the MD/RS model for NDUFB9, and the MD/Iso and MD/RS models for CLASP1 and YWHAZ. Protein levels were normalized to the mean of the controls. Individual values represent the protein levels for each subject or animal. Means and standard deviation are shown in the graphs. ns: not significant, *p<0.05, **p<0.01, ***p<0.001, ****p<0.0001.

## DISCUSSION

Our study identified and characterized the proteomic alterations for the cerebellar cortex in chronic schizophrenia. To the best of our knowledge, this is the first time in which the individual proteomic signature allows to segregate most SZ cases from healthy controls in unsupervised analysis, similarly as achieved in some gene expression studies [27,43]. A subset of the molecular findings of this study had been previously related to brain diseases, the main ones being mitochondrial and mental disorders, but also other less represented diseases such as stress. In this context, a study related psychosocial stress mediated by the cortisol with decreased blood flow in the cerebellum and an impairment of retrieval memory [44]. This evidence suggests that the cerebellum is a brain area susceptible to psychosocial stress. This susceptibility could be exacerbated in SZ, thereby contributing to the cognitive symptoms in this disorder. Here we also observed an enrichment of proteins involved in neurodegenerative diseases. Although the neurodegenerative hypothesis is strongly debated in SZ [45,46], enrichment in the neurodegenerative disease category in this study could be related to the up-regulation of apoptosis, a biological process also found enriched in this work. In this study, the enrichment of the mitochondrial disease category in the altered proteins could be related to a down-regulation of biological processes related to oxidative phosphorylation and TCA, biological processes that were also observed to be enriched in this study. In SZ, altered mitochondrial function has been previously reported [47].

Furthermore, our network analyses showed two modules for the down-regulated network, namely energy metabolism and neutrophil degranulation. For the up-regulated network, we found pathway crosstalk between proteins of vesicle-mediated transport, axon guidance, signalling by Rho GTPases, apoptosis, and the cell cycle, suggesting that signalling pathways are dynamic events in which the alteration of one protein could affect several pathways.

### Energy metabolism module

We found a down-regulation of proteins involved in the energy production in the cerebellum in schizophrenia. The cerebellum represents 11% human brain weight [48] and the distribution of the energy it uses varies between different cell types. The more energy-demanding functions in Purkinje cells include the production of action potentials and maintenance of postsynaptic receptors, while granule cells consume more energy to propagate action potentials and support the resting potential [49]. In the context of SZ, a study of cerebellar activity showed decreased blood flow in this area during several tasks, including attention, social cognition, emotion and memory [22,50]. Evidence of mitochondrial dysfunction in SZ includes genetic [51], metabolic [52], and enzymatic dysfunctions [53], anatomical abnormalities[54], and disturbed levels of proteins of glycolysis, TCA cycle, mitochondrial function and oxidative stress [55–58]. These studies are in line with our results showing a down-regulation of energy network built by TCA cycle/respiratory electron transport and several mitochondrial proteins. In our study, NDUFB9 was observed to be one of the down-regulated proteins in SZ involved in respiratory electron transport. NDUFB9 is widely expressed in the cerebellum with moderate expression in cerebellar granule layer and Purkinje layer. It was also observed to be down-regulated in the double-hit MD/Iso murine model but not in the MD/RS model. NDUFB9 is involved in the assembly of Complex I. Together with the activity of Complex I, NDUFB9 has been investigated in other brain areas in SZ [59,60]. Thus, the altered expression of NDUFB9 observed in this study could contribute to reduce energy metabolism in cerebellar cells through the disruption of Complex I in respiratory electron transport and could consequently decrease the propagation action potentials among the major cell types of the cerebellar cortex in chronic SZ.

### Neutrophil degranulation module

Our study observed down-regulation of several proteins involved in the various processes of the neutrophil degranulation pathway in cerebellar tissue. Neutrophil degranulation is one of the first defence barriers against infection [61]. An imbalance of the immune system is one of the hypotheses underlying SZ [62]. The altered proteins of neutrophil degranulation were related to processes such as transmigration, sequestration of microbes in the autophagosome, and exocytosis of primary and second granules. In our study, METTL7A was observed to be consistently down-regulated in both SZ human samples and our SZ murine models. METTL7A, a member of the METTL family of methyltransferases, is of interest as it has been poorly studied; indeed, only one study investigated its role in RhoBTB1 signalling in maintaining Golgi integrity [63], a function that could be involved in neutrophil degranulation. In the context of neutrophil degranulation, the altered expression of METTL7A could impair Golgi integrity and contribute to the abnormal formation of secretory granules in neutrophils, altering the first defence barrier of the innate immune response in chronic SZ. Limited information is available about METTL7A deregulation in the context of schizophrenia. Previous studies have reported a decrease in the RNA and protein levels on the prefrontal cortex (Brodmann area 46/10) and the anterior cingulate cortex (Brodmann area 24) in SZ subjects, respectively [64,65]. Another study on the prefrontal cortex (Brodmann area 9) reported an increase in the RNA levels of METTL7A [66]. However, its expression profile in the brain and its function in SZ are completely unknown and further studies are needed for METTL7A function in brain.

### Vesicle-mediated transport pathways

In our proteomic study we found up-regulated pathways related to vesicle-mediated transport. Transport is a critical step for the correct maintenance of biological processes. In neurons, altered transport could limit effectiveness in neuronal communication [67]. Defective synaptic transmission and neurotransmitter release [68], decreased in pre-synaptic vesicle proteins [69,70], and altered levels of proteins involved in synaptic vesicle fusion [71,72] have been associated with SZ. However, in our study we found an increase of this pathway that could be indicative of a compensatory strategy. YWHAZ was found to be up-regulated in our proteomic study. We also studied the effect of two double-hit SZ murine models on YWHAZ. However, we did not observe any significant changes in YWHAZ in these stress-based models. Other proteomics studies in different brain regions of SZ subjects have shown down-regulation of YWHAZ [73,74]. The role of YWHAZ as an adaptor protein of extracellular vesicles (EVs) involves the stabilization of vesicles and synapsis [75]. The overexpression of YWHAZ could increase the formation and release of EVs carrying protein or miRNA to the synapsis, which could be a compensatory mechanism for a defective synaptic activity in this disorder in the cerebellum.

### Axon guidance pathway

Our proteomic study showed an altered axon guidance pathway in the cerebellum. Defects in neuronal connectivity during development have been proposed as an important cause of the etiopathology of SZ [76,77]. A study based on the SZ-GWAS database found the axon guidance pathway to be altered in this disorder [78]. CLASP1 was a protein from axon guidance down-regulated in schizophrenia and further studied in two murine stress models in this study. Our result showed decreased expression of CLASP1 in the MD/Iso model, while our proteomic study in SZ subjects showed an up-regulation. This protein participates in neurite outgrowth by binding at microtubules [79]. One possible explanation for the up-regulation of CLASP1 in human cerebellum could be the accumulation of this protein in the neuritic growth cone due to a decrease in energy production required for the assembly of CLASP1 at microtubules.

The use of human *postmortem* brain constitutes a valuable tool to understand the molecular pathways altered in several psychiatric disorders. However, it has limitations. Firstly, confounding factors such as age, *postmortem* delay, and pH have to be carefully explored. In our proteomic study, we did not find any association between these variables and the significantly altered proteins in the cerebellum. Secondly, patients with chronic schizophrenia had long-term and heterogeneous antipsychotics medications. Thirdly, our study only included men, who do not represent a real population of this disorder. Fourthly, our study consisted entirely of elderly individuals due to the type of sample available.

In summary, our study reveals altered energy, immune and axonal-related networks in the cerebellum in schizophrenia, suggesting that the accumulation of altered events some related to stress in these networks could lead to a dysfunction of the molecular network in the cerebellum in chronic schizophrenia impacting on normal function of the cerebellar circuits. Further the validation of the most robust candidates of each networks in two double hit murine models for schizophrenia point out that some of the molecular network alterations observed in schizophrenia could be induced by different combinations of stress exposure during the development of the cerebellum. Thus, our results suggest that the cerebellum is an organ that accumulate molecular errors during early postnatal life, could be susceptible to psychosocial stress and that this vulnerability could be increased in schizophrenia. We believe these findings could provide possible molecular targets to explore treatments for schizophrenia in future studies. Moreover, further studies in the cerebellum that deep in altered networks related to stress during postnatal maturation are needed in the context of schizophrenia.

## Acknowledgement

The authors thank the donors and their families for the donation of their brains; the collaboration of the team of the Hospital Universitari de Bellvitge Brain Bank, and the team of the Banc de Teixits Neurologics of Parc Sanitari Sant Joan de Déu for their help. We thank Èlia Vila, Beatriz Moreno and Álvaro González-Bris for her technical support and Dr Rose for the English editing of this manuscript.

## Author contributions

**Author A.V:** acquired data, interpreted the results and co-wrote the first draft of the manuscript. **Author R.R-M:** designed, conducted and analysed the proteomic experiments and contributed to the discussion of the results. **Author K.M:** developed the double-hit animal models of SZ and acquired data from them and contributed to the discussion of the results. **Author B.G.B:** developed the double-hit animal models of SZ and contributed to the discussion of the results. **Author J.C.L:** contributed to the discussion of the results. **Author J.V:** supervised the proteomic analyses and contributed to the discussion of the results. **Author B.R:** designed the study, contributed to the interpretation and discussion of the results, and co-wrote the first draft of the manuscript. All authors revised the manuscript critically and approved the final article.

## Supplementary Material

Disease classification was performed using the PharmMGKB database and genes associated with individual disease terms were inferred using GLAD4U. For the gene ontology analysis we used the Gene Ontology Consortium database: pathways were identified according to the Reactome database and proteins assigned to each pathway are listed. Total number: number of reference proteins in category/pathways; Observed number: proteins in the data set and also in category/pathways; E: expected in the category and adjusted p-value is corrected for test multiple, FDR=0.1. These analyses were carried out using Webgestalt.

