## Supplementary Material for "Analysis Of Molecular Networks In The Cerebellum In Chronic Schizophrenia: Modulation By Early Postnatal Life Stressors In Murine Models"

### Supplementary materials and methods

##### Protein reduction, alkylation, LysC digestion, and desalting

Protein extracts were prepared from tissue samples using NP40 lysis buffer as described previously [33]. Protein concentration was determined by Bradford assay (Biorad, Hercules, CA, USA). 200 µg of total protein was lyophilized (Telstar, Lyoquest-55) per sample for mass spectrometry analysis. 200 µg of total protein extracts lyophilized from control and SZ lysates were resuspended in 100 µl 8M Urea, 50 mM Tris pH 8.2, 100 mM NaCl. The lysate was reduced with 3 mM dithiothreitol at 55 ºC for 30 min, alkylated with 20 mM iodoacetamide at room temperature for 30 min and quenched with additional 3mM dithiothreitol for 30 min at room temperature. The extract was digested with the endoproteinase LysC at 10 ng/µl at 30ºC overnight. Peptides were desalting with OASIS MCX cartridge (Waters): cartridge was conditioned with Methanol, 5% ammonium hydroxide in Methanol, and 0.1% TFA; acidified peptides were loaded on the column and washed with TFA 0.1% and 0.1% Formic acid in Methanol, finally peptides were eluted using 5% ammonium hydroxide in Methanol. Eluted peptides were acidified with formic acid. and dried before re-suspending in MS Buffer composed of 3% formic acid, and 4 % acetonitrile in water.

##### Liquid chromatography coupled to tandem mass spectrometry

The LC/MS-MS analysis was performed in a Q-Exactive mass spectrometer( Thermofisher Scientific, CA, USA) coupled to an Easy nLC II liquid chromatography system. Peptides were loaded onto a 100 µm ID × 3 cm precolumn packed with Reprosil C18 3 µm beads (Dr. Maisch GmbH), and separated by reverse-phase chromatography on a 100 µm ID × 30 cm analytical column packed with Reprosil C18 1.9 µm beads (Dr. Maisch GmbH). A gradient of 3% to 30% acetonitrile in 0.125% formic acid was used delivered at 225 nl/min over 130 minutes, with a total 180-minute acquisition time. Peptides were analyzed online on the orbitrap mass analyser using a top 20 data-dependent acquisition with all MS spectra being acquired and stored in centroid mode. Full MS scans were acquired from 300 to 1500 m/z at 70,000 FWHM resolution with a fill target of 3E6 ions and maximum injection time of 100 ms. The 20 most abundant ions on the full MS scan were selected for fragmentation using 2 m/z precursor isolation window and beam-type collisional-activation dissociation (HCD) with 26% normalized collision energy. MS/MS spectra were collected at 17,500 FWHM resolution with a fill target of 5E4 ions and maximum injection time of 50 ms. Fragmented precursors were dynamically excluded from selection for 35 s.

**Data analysis**

Raw files were processed and analysed using MaxQuant (version 1.5.3.8). MS/MS spectra were searched with Andromeda against UniProt fasta UP000005640 (Downloaded: 2015-06-30) with common contaminants added. The precursor mass tolerance was set to 7 ppm, and the fragment ion tolerance was set to 20 ppm. Search parameters included full LysC enzyme specificity with up to three missed cleavages permitted. The target-decoy database search strategy was used to guide filtering and estimate false discovery rates (FDR). Peptides matches were filtered to an FDR of ≤0.01. Proteins with at least one peptide were considered identified. Label-free quantification (LFQ) was selected for individual protein comparisons between control and schizophrenia groups. A quality cut-off for protein determination was the presence of the protein in at least 7 samples per group. The normalized LFQ intensity was referred to media of the controls. A significance value for each quantified protein was calculated using Student's t-test and the correction of significance values of the quantified protein data set was performed following the Benjamini and Hochberg methods [34]. An FDR was computed for all significant values and the FDR threshold was set to 0.1.

The quantified proteins were imported into Perseus software platform (version 1.6.1.3) to check data quality obtained by LC/MS/MS and to visualize the data distribution [35]. We performed the following analyses. 1. Correlation matrix: the normalized LFQ intensity data were used to estimate a correlation coefficient matrix between controls and schizophrenia patients. For easy visualization, the correlation coefficient data were plotted as a heat map. 2. Hierarchical Clustering: This was carried out on Z-score transformed normalized LFQ intensity data for each protein using Euclidean distance.

**Murine model for schizophrenia**

Three pregnant Wistar rats (Harlan Ibérica, Spain) at gestation days 14-16 were individually housed in a temperature/humidity-controlled environment in a 12h light/dark cycle with free access to food and water. The litter sizes varied between 7 and 11 animals. One was used as a control group and the others as part of a double-hit model. After birth, at postnatal day 9 (PD9), both double-hit litters were exposed to maternal deprivation for 24hrs as a first hit; this early stressful life event has an impact on prepulse inhibition in rats, similar to the alterations seen in schizophrenic patients. On PD21, the pups were weaned and one of the litters was exposed to isolation for 5 weeks as a second hit (PD21-56); although the pups were housed separately (1/cage), they could smell, hear and see their siblings, but they could not come into physical contact with them. Following isolation, the animals were regrouped (DM/Iso). Meanwhile, the second hit for the other litter exposed to maternal deprivation involved restraint stress between PD 72 to 78 for 6 hours every day (DM/RS). These conditions represent suitable rodent models for the study of neuropsychiatric dysfunctions (Bailoo *et al.*, 2016; van Zyl, Dimatelis and Russell, 2016). Group sample sizes were CT, n=11; DM/Iso, n=9; DM/RS, n=7. All experimental protocols adhered to the guidelines of the Animal Welfare Committee of the Complutense University in accordance with European legislation (D2010/63/UE). All efforts were made to reduce the number of animals used and minimize animal suffering in the experiments.

**Western Blot analysis**

Brain cerebellum samples were homogenized by sonication in PBS (pH=7) mixed with a protease inhibitor cocktail (Complete®, Roche, Spain). After determining and adjusting protein levels, homogenates of cerebellum tissue were mixed with Laemmli sample buffer (Bio-Rad, USA) and β-mercaptoethanol (50 µl/ml Laemmli), 15µg were loaded into an electrophoresis gel. Once separated on the basis of molecular weight, proteins from the gels were blotted onto a nitrocellulose membrane with a semi-dry transfer system (Bio-Rad) and were incubated with specific antibodies against: (1) Methyltransferase-like protein 7A (METTL7A, 1:750 in BSA 1%; ABclonal A8201); (2) NADH dehydrogenase (ubiquinone) 1 beta subcomplex, 9 (NDUFB9, 1:1000 in BSA 0,5%; sc398869, SCT); (3) cytoplasmic linker associated protein 1 (CLASP1, 1:1000 in BSA 0,5%; sc390159, SCT); (4) tyrosine 3-monooxygenase/tryptophan 5-monooxygenase activation protein zeta (YWHAZ, 1:1000 in BSA 0,5%; sc293415, SCT); (5) β-actin (1:10000; A5441, Sigma). Primary antibodies were recognized by the respective horseradish peroxidase-linked secondary antibodies. Blots were imaged using an Odyssey Fc System (Li-COR, Biosciences, Germany) and were quantified by densitometry (NIH ImageJ software). In all the WB analyses, the housekeeping protein β actin was used as a loading control, and each western blot was performed at least three times in separate assays. The data were presented as fold change from the control group.

#### Table S1: Demographic, clinical and tissue-related features of cases.

| | Schizophrenia (n=12) | Control (n=14) | Statistic | *p*- value |
|---|---|---|---|---|
| **Gender** | | | | |
| Male | 100% (*n*=12) | 100% (*n*=14) | N/A | N/A |
| Age (years) | 72 ± 9 | 69 ± 11 | 78.5[a] | 0.56 |
| PMD (hours) | 5.48 ± 2.29 | 5.46 ± 1.81 | 0.02; 25 [b] | 0.98 |
| pH Cerebellum | 6.88 ± 0.49 | 6.61 ± 0.63 | 1.25; 25 [b] | 0.22 |
| **SZ diagnosis** | | | | |
| Chronic residual | 66.67% (n= 8) | | | |
| chronic paranoid | 16.67% ( n=2) | | | |
| chronic disorganized | 8.33% (n=1) | | | |
| chronic catatonic | 8.33% (n= 1) | | | |
| Age of onset of SZ (years) | 22 ± 8 | N/A | N/A | N/A |
| Duration of illness | 50 ± 9 | N/A | N/A | N/A |
| **Toxicology** | | | | |
| Daily AP dose (mg/day) [c] | 609 ± 507.10 | N/A | N/A | N/A |
| First generation AP | 16.67% (n=2) | | | |
| Second generation AP | 58.33% (n=7) | | | |
| First and Second generation AP | 8.88% (n=1) | | | |
| AP free | 16.67% (n=2) | | | |

Mean ± standard deviation; PMD, postmortem delay; SZ, schizophrenia; AP, antipsychotics; N/A, not applicable.
a Mann-Whitney U for non-parametric variables.
b T-statistic and degree of freedom for parametric variables.
c Last daily chlorpromazine equivalent dose was calculated based on the electronic records of drugs prescriptions of the patients as described (Gardner et al., 2010).

**Table S2:** Proteins previously reported in proteomic studies in schizophrenia

| Cerebral area | Proteins previously reported in schizophrenia | References |
|---|---|---|
| **Cerebellum** | CAMK4; C11orf58; NDUFA12; AHCY; ERP44; PAFAH1B1; PRDX6; NCKAP1; LCP1; MYH9; NAXE; ACLY; CYB5R1; NAPA; CCT7; HSD17B4; BCL2L13; RAB21; RIMS1; UBE2I | Vidal-Domènech et al., 2020; Reis-de-Oliveira et al., 2020 |
| **Anterior cingulade** | CS; GPD1; AHCY; CBR1; UCHL1; PGAM1 | Clark et al., 2007 |
| **Anterior cingulade cortex** | CS; ACAT2; SOD1;DDAH1;DCTN2; IDH2; HNRNPK; IARS; METTL7A; ACTR2; PPIA; LIN7A; PIP4K2A; NAPA; SPTAN1; SCRN1; TPM1; ARHGDIA; YWHAZ; YWHAB; COMT; HSD17B4; AP2M1; UCHL1 | Clark et al., 2007; ClarK et al., 2006; Föcking et al., 2015; Martins de Souza et al., 2010; Smalla et al., 2008; English et al., 2011 |
| **Anterior hippocampus** | UCHL1 ; DDAH1 ; SOD1 ; CAPZA2 ; YWHAE; YWHAZ ;PRDX6 | Nesvaderani et al., 2009 |
| **Posterior hippocampus** | PGAM1 | Nesvaderani et al., 2009 |
| **Anterior temporal lobe** | YWHAQ; YWHAZ; HSPD1; NDUFB5; SPTAN1; YWHAH; YWHAE; MYH9; PURA; HNRNPK; MOG | Saia-Cereida et al., 2017; Martins de Souza et al., 2009[b] |
| **Auditory cortex** | GNAQ; ATP1A3; UCHL1 | Mc Donald et al., 2015 |
| **Corpus callosum** | YWHAZ; UCHL1; MOG; NPM1; DDAH1; YWHAE; YWHAB; ANK3; SOD1; DDT; HP1BP3; HNRNPR; YWHAH | English et al., 2011; Saia-Cereida et al., 2015; Sivagnanasundaram et al., 2007; Saia-Cereida et al., 2016 |
| **Dorsolateral prefrontal cortex** | DPYSL5; NDUFS1; HSPD1; SPTAN1; YWHAE; PGAM2; MOG; RAP2A; NDUFA12; GDAP1; NDUFB10; SIRPA; HP1BP3; DPYSL4; ADH5; GSTM3; IGSF8; CLSTN1; MAP6; PRDX6; TKT; ERP29; PGAM1; PURA; YWHAZ; CBR1; TCP1; HSPA9; BCL2L13; UCHL1; ARPC1A; ITGAV; SCRN1; GPM6A; MARCKS; TXN2; TPT1; CYB5A; RTN1; GNAQ; ATP5PD; PPA2 | Saia-Cereida et al., 2015; Martins de Souza et al., 2009[a]; English et al., 2009; Martins de Souza et al., 2009[c]; Chan et al., 2011; Behan et al., 2009; Pinacho et al., 2016; Pennington et al., 2008; English et al., 2011; Wesseling et al., 2013; Novikova et al., 2006; Smalla et al., 2008; Prabakaran et al., 2004 |
| **Hippocampus** | TPPP; UBE2M; YWHAE;SOD1; PRDX6; TCP1; UCHL1 | Schubert et al., 2015; Föcking et al., 2011 |
| **Mediodorsal thalamus** | YWHAZ; CBR1; SYN3; PGAM1; NDUFB9; TKT; MOG; HSPD1; PPIA; PITPNA | Martins de Souza et al., 2010 |
| **Orbitofrontal prefrontal cortex** | RAB35; ATP2B3; PGAM1; RTN1; DDAH1; ATP1A1; MARCKS; MOG; YWHAH; SYN3; GPM6A; ATP1A3 | Velasquez et al., 2017 |
| **Wernicke's area** | DLD; PRDX6; PGAM1; NDUFS1 | Martins de Souza et al., 2009[d] |
| **Insular cortex** | SNCG; ERLIN2; NPM1; COPS4 | Pennington et al., 2008 |

**Figure S1**

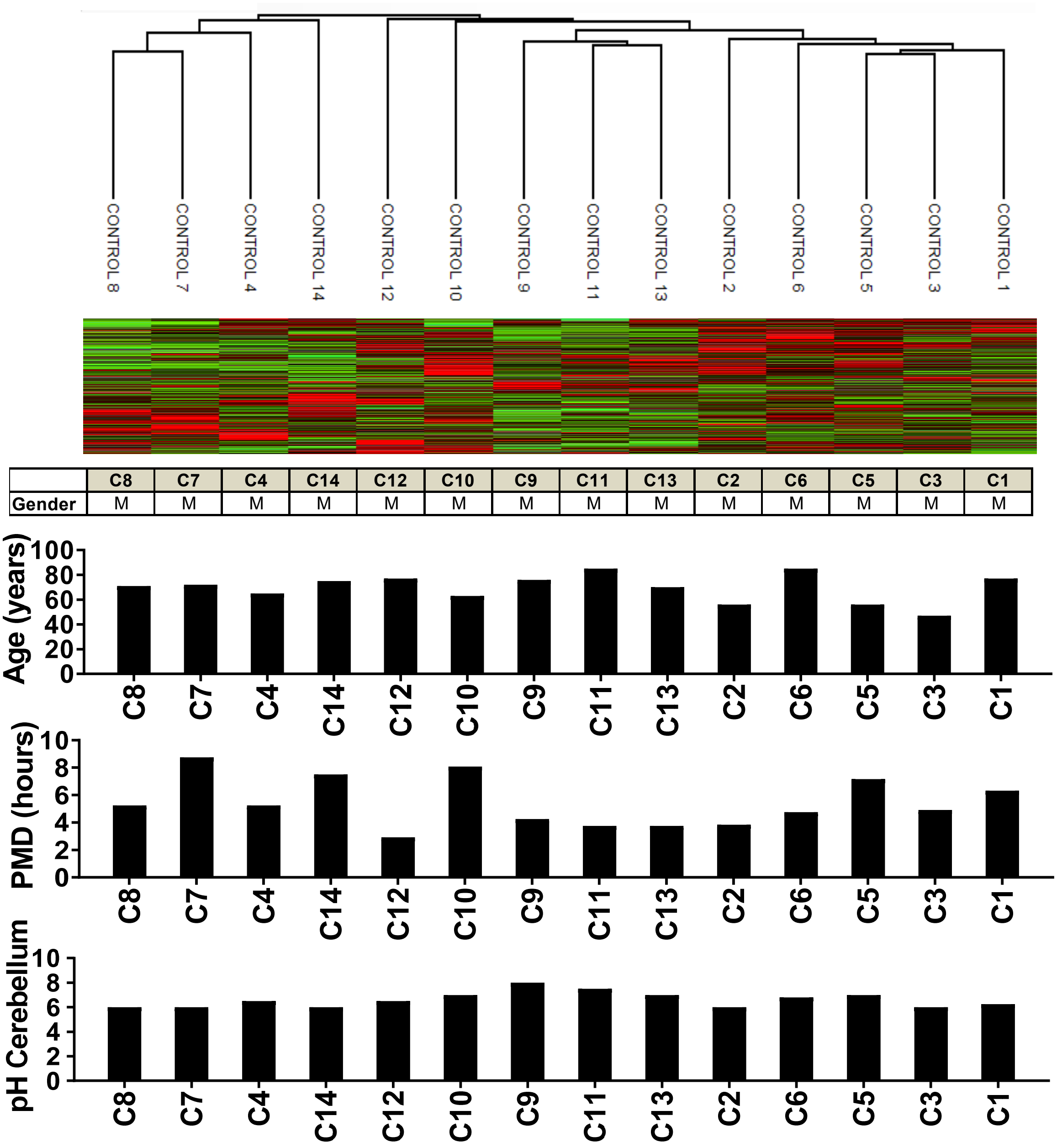


**Figure S1:** Unsupervised hierarchical clustering analysis for quantified proteins from *postmortem* cerebellum of 14 healthy control samples. The table below it shows the gender. The graphs show the tissue-related features of each healthy control sample. M: Male, PMD: *Postmortem* time delay.

#### Figure S2

**A. TCA-Respiratory electron transport (NDUFB9, MW: 22 kDa; MW: ß-actin 42 kDa)**

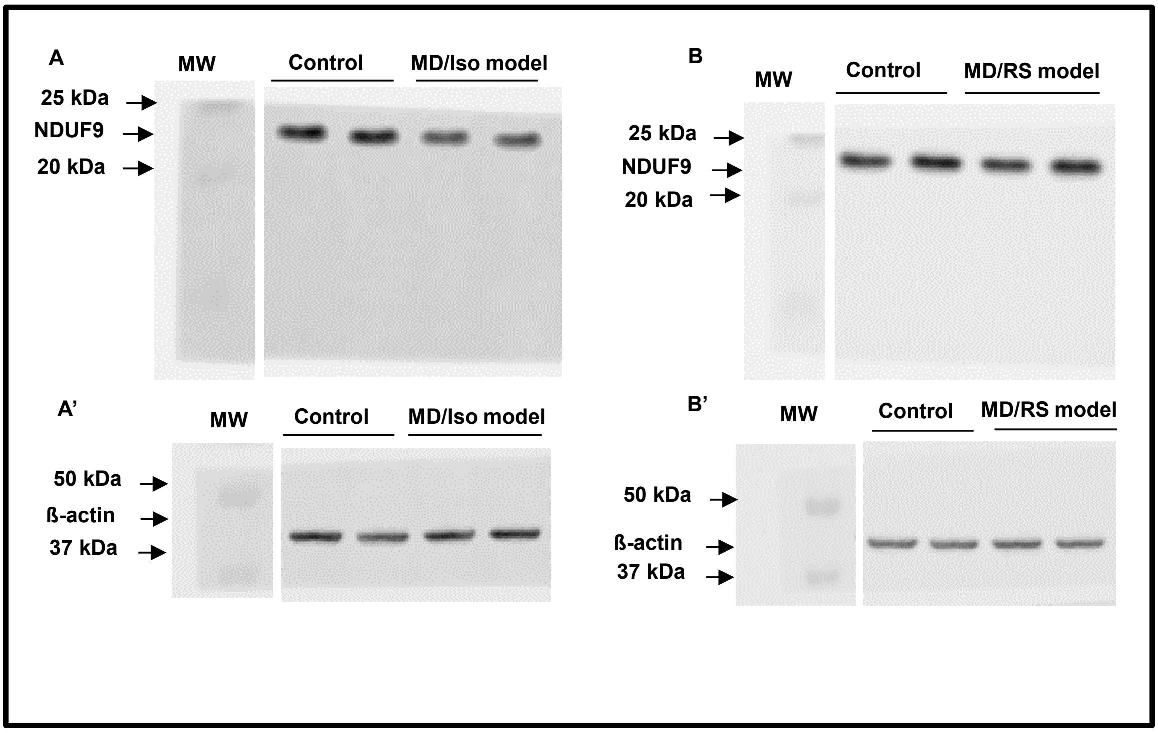


**B. Neutrophil degranulation (METTL7A, MW: 28 kDa; MW: ß-actin 42 kDa)**

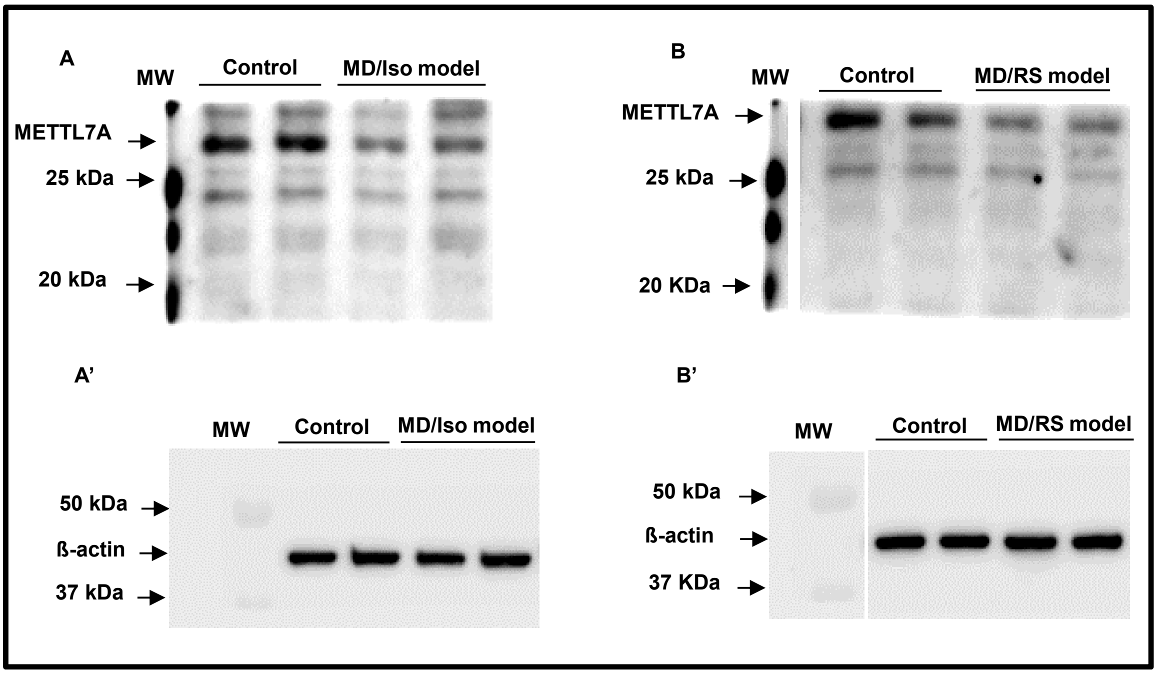

#### Continuation of Figure S2

C. Axonal guidance (CLASP1, MW: 160 kDa; MW: ß-actin 42 kDa)

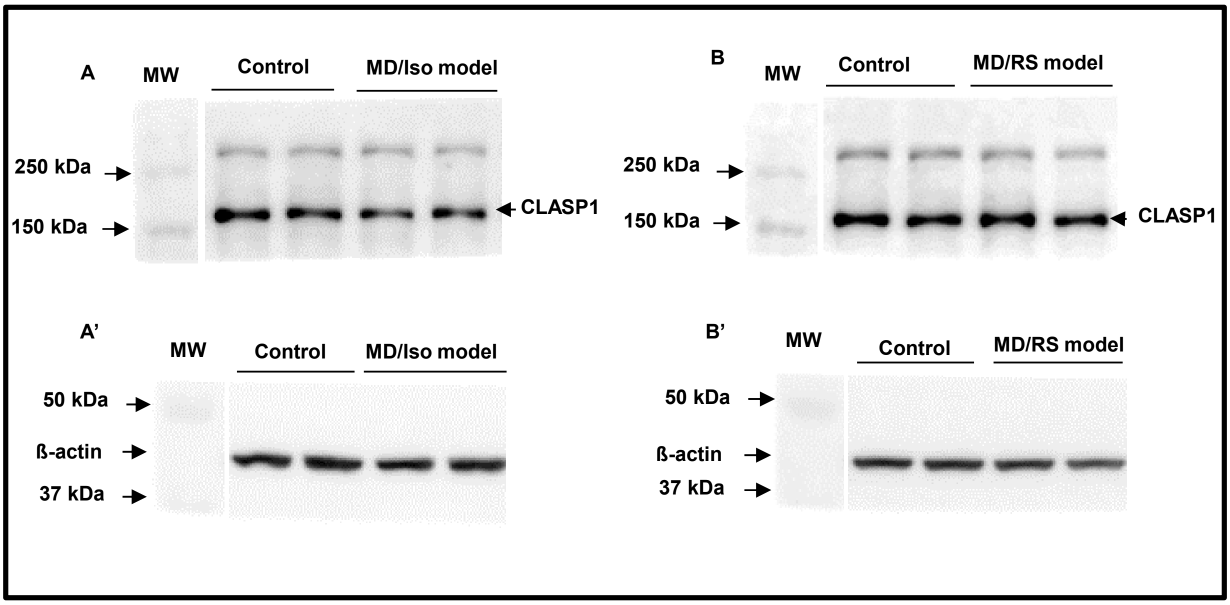


D. Vesicle-mediated transport (YWHAZ, MW: 28 kDa; MW: ß-actin 42 kDa)

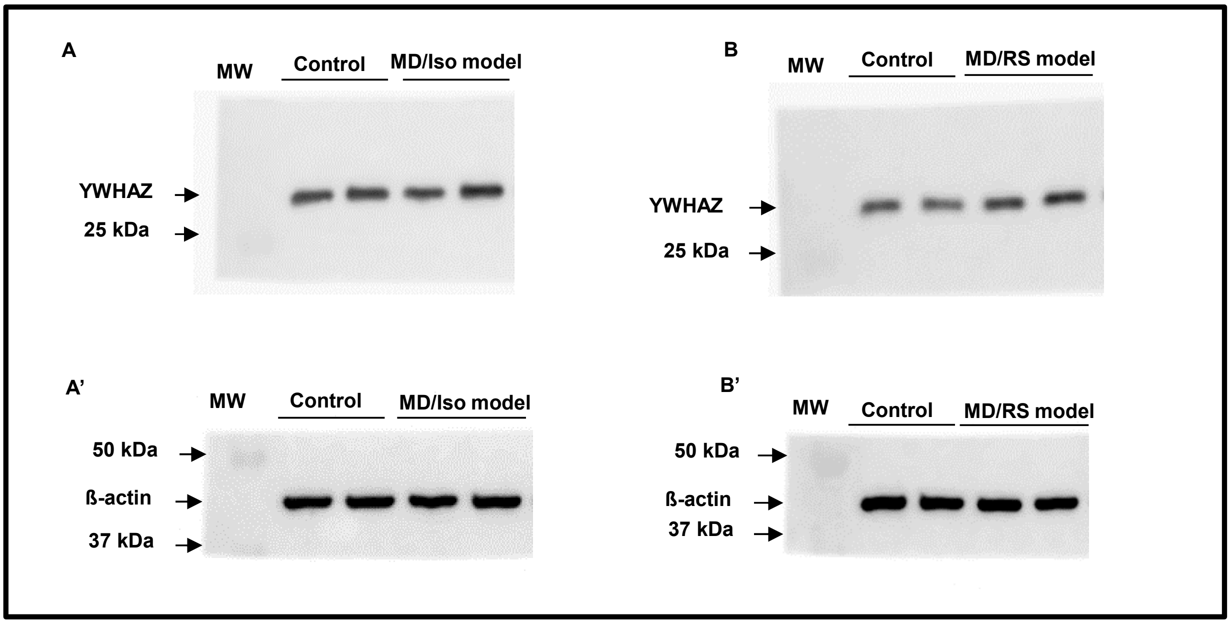

**Figure S2. Validation of hit proteins in two double-hit murine models for schizophrenia.** The hit proteins of the enriched pathways in our proteomic study were analysed in two double-hit murine models for SZ**. Panel A**. Immunoblot of NDUFB9, a hit protein from TCA/Respiratory electron transport. **A**. Reduced expression of NDUFB9 in the MD/Iso model in comparison with controls. **B**. Expression of NDUFB9 in the MD/RS model and controls. No significant changes were found in this model. **A**' and **B'** show ß-actin expression used as a loading control. **Panel B**. Immunoblot of METTL7A, a hit protein from neutrophil degranulation. **A**. Immunodetection of METTL7A in the MD/Iso model in which it was significantly downregulated in comparison with the control. **B**. Expression of METTL7A in the MD/RS model in which it was also significantly downregulated with respect to the control. **A**' and **B'** show ß-actin expression used as a loading control for METTL7A. **Panel C**. Immunoblot of CLASP1, a hit protein from axon guidance. **A**. Expression of CLASP1 in the MD/Iso model in which it was significantly decreased in comparison with the control. **B**. Expression of CLASP1 in the MD/RS model; no significant changes were observed in this model. **A**' and **B'** show ß-actin expression used as a loading control. **Panel D**. Immunodetection of YWHAZ a hit protein from vesicle-mediated transport. In this analysis we did not observe any change in YWHAZ expression either in the MD/Iso or the MD/RS model in comparison with controls. **A**' and **B'** show ß-Actin expression used as a loading control. MD/Iso: maternal deprivation and isolation model. MD/RS: Maternal deprivation and restraint stress model. MW: Molecular weight. All protein levels were normalized to ß-actin values.
